## Supplementary Perceptual Masking Demonstration for "Neural Correlates of Perceptual Similarity Masking in Primate V1"

### Stimulus Sheets Demonstrating Orientation and Spatial Frequency Masking

The following four pairs of slides demonstrate orientation and spatial frequency masking. To view the slides, it is probably best to use full screen mode (cntl L in Adobe Acrobat or Reader). The first slide in each pair shows an array of raised-cosine-windowed sinewave targets of a given amplitude (in units of gray level), added to a uniform background with gray level 128 (out of 256). The range of orientations (in degrees) is indicated on the vertical axis and the range of spatial frequencies (in cycles/degree) is indicated on the horizontal axis. The spatial frequencies assume a particular viewing distance. This viewing distance can be computed by measuring the width of the sheet (the gray area) on your display and then dividing that width by 0.1223, which gives a sheet subtending 7 deg, with each target subtending 1 deg. The second slide in each pair shows the effect of adding a vertical sinewave grating of 4 cycles/degree and a mean-to-peak amplitude of 64 (Michelson contrast of 0.5).

Toggling the masking grating off and on while fixating a specific target location shows the masking effect in the fovea for the specific target at that location.  Fixating at various locations and attending to the target visibility at all the locations gives some appreciation for the effects of retinal location on detectability for different orientations and spatial frequencies.

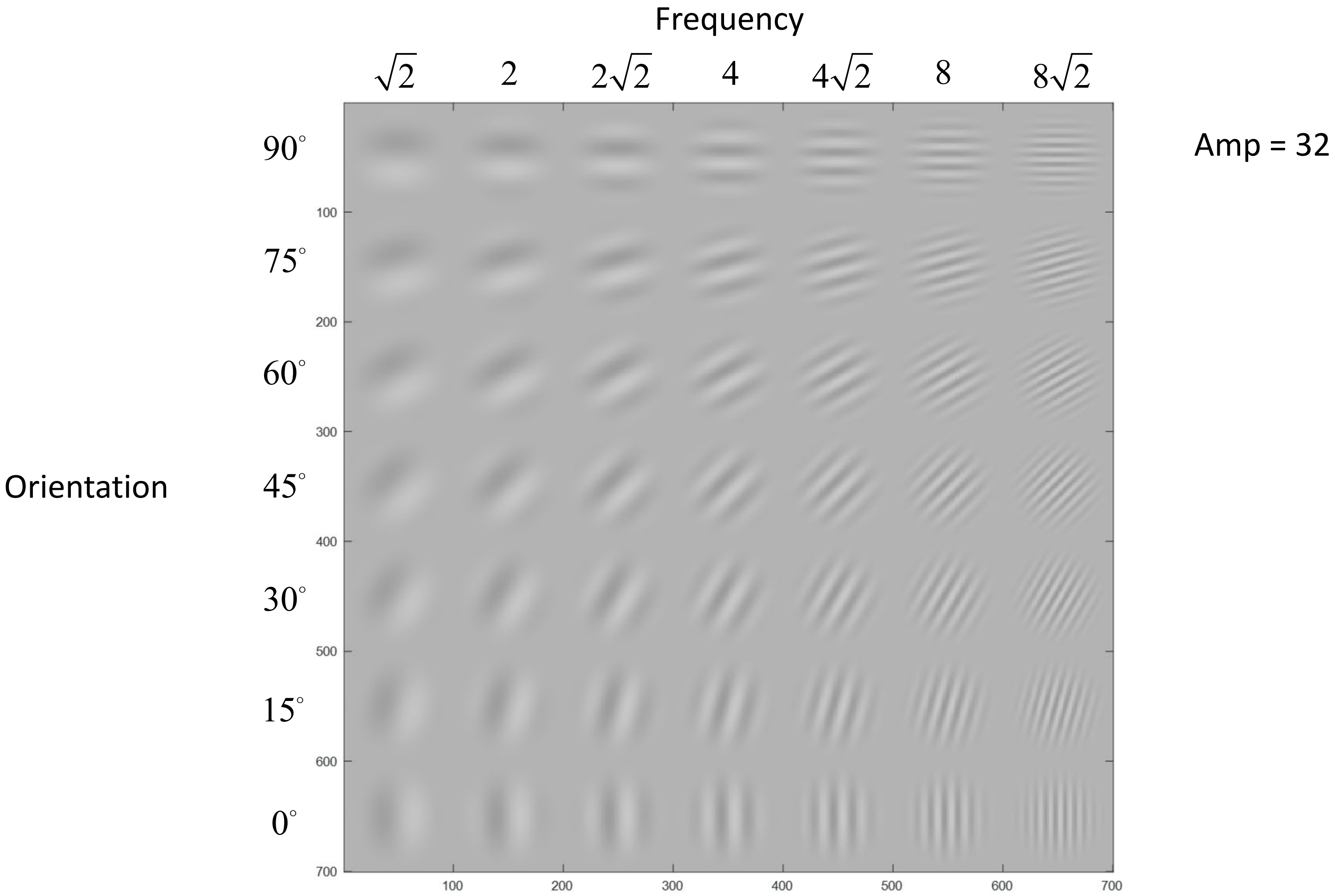
Frequency
$\sqrt{2}$
2
$2\sqrt{2}$
4
$4\sqrt{2}$
8
$8\sqrt{2}$
Amp = 32
Orientation
90°
75°
60°
45°
30°
15°
0°
100
200
300
400
500
600
700

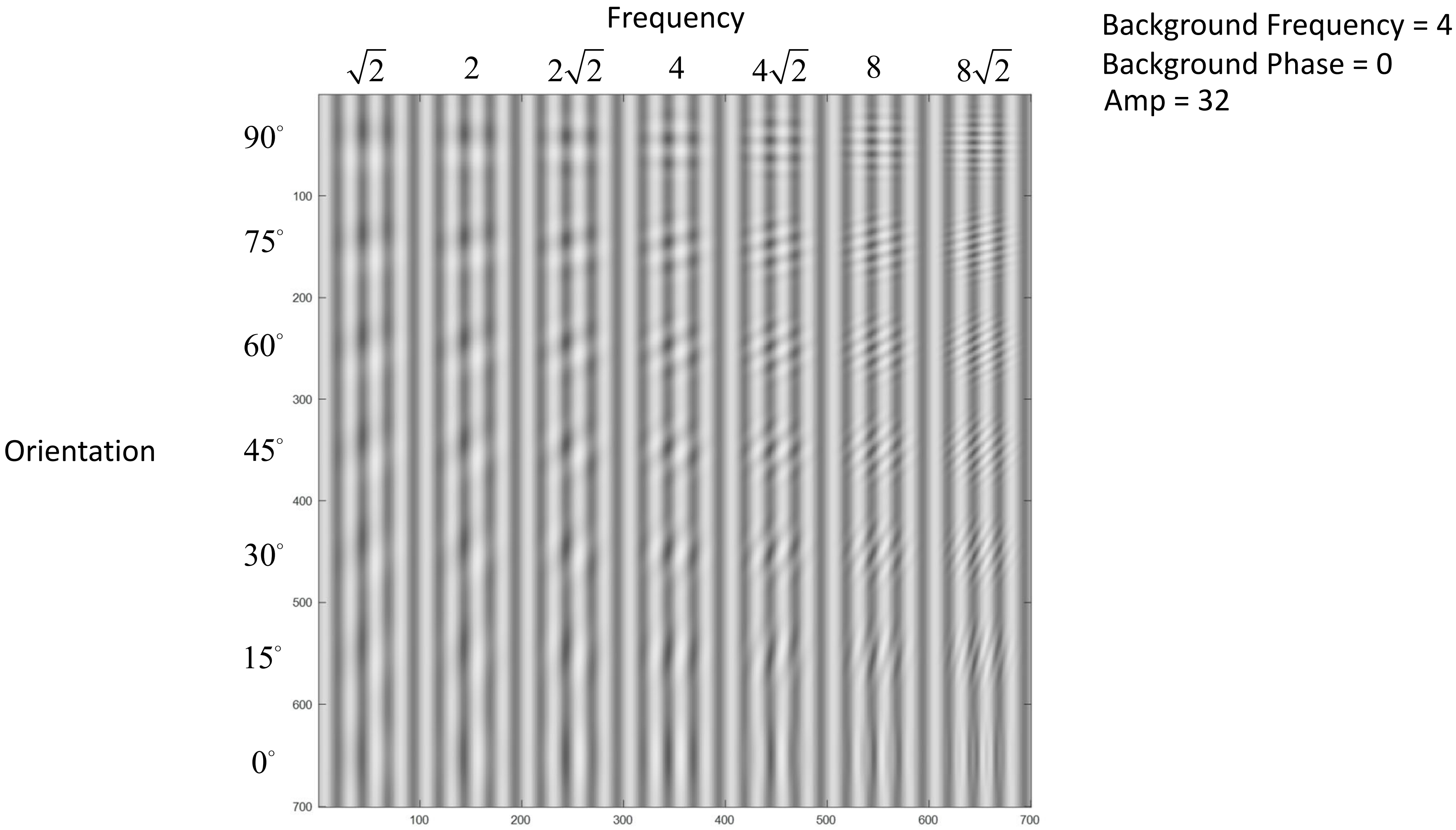
Frequency
$\sqrt{2}$
2
$2\sqrt{2}$
4
$4\sqrt{2}$
8
$8\sqrt{2}$
Orientation
90°
75°
60°
45°
30°
15°
0°
Background Frequency = 4
Background Phase = 0
Amp = 32

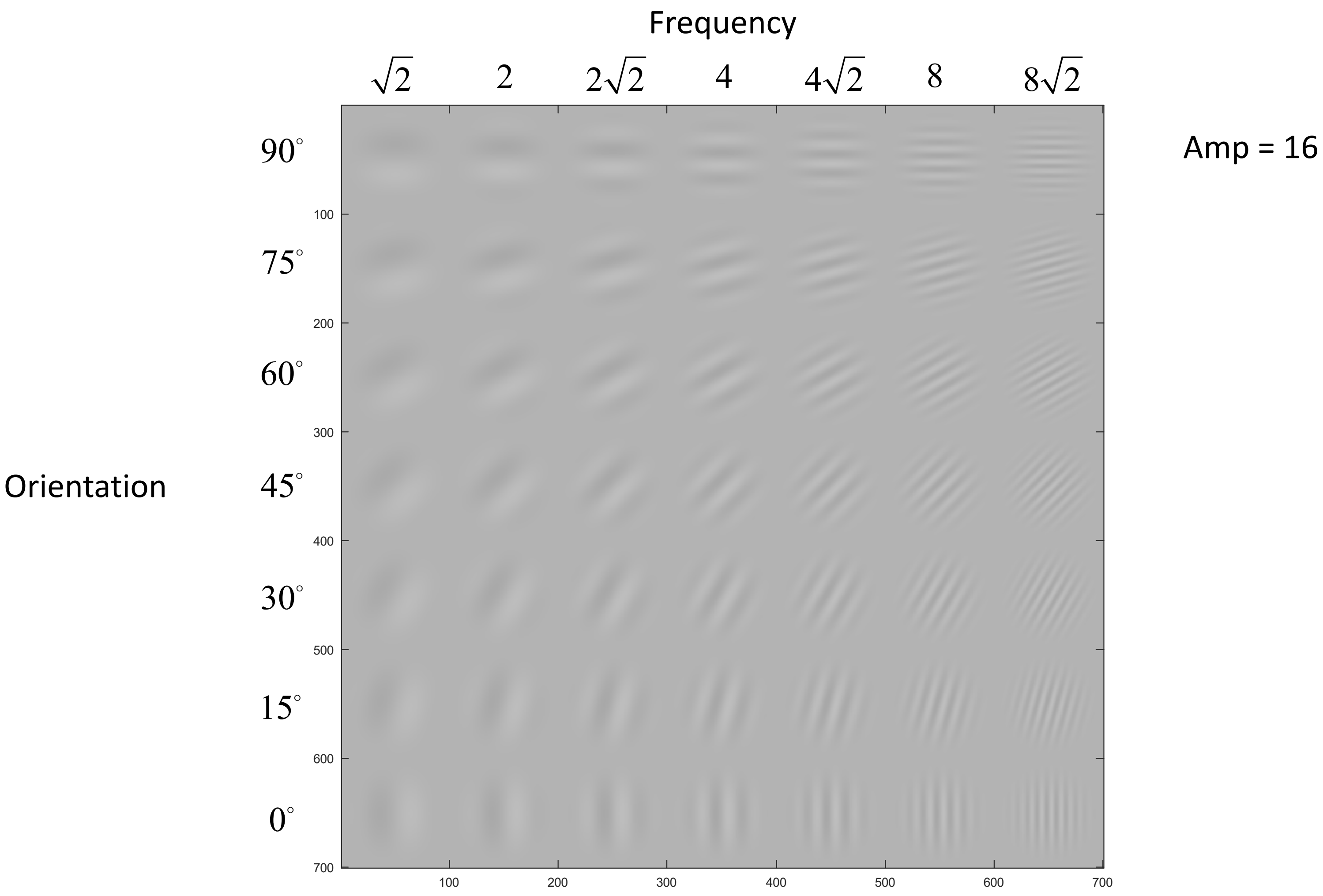

Frequency
$\sqrt{2}$
2
$2\sqrt{2}$
4
$4\sqrt{2}$
8
$8\sqrt{2}$
Amp = 16
Orientation
90°
75°
60°
45°
30°
15°
0°
100
200
300
400
500
600
700

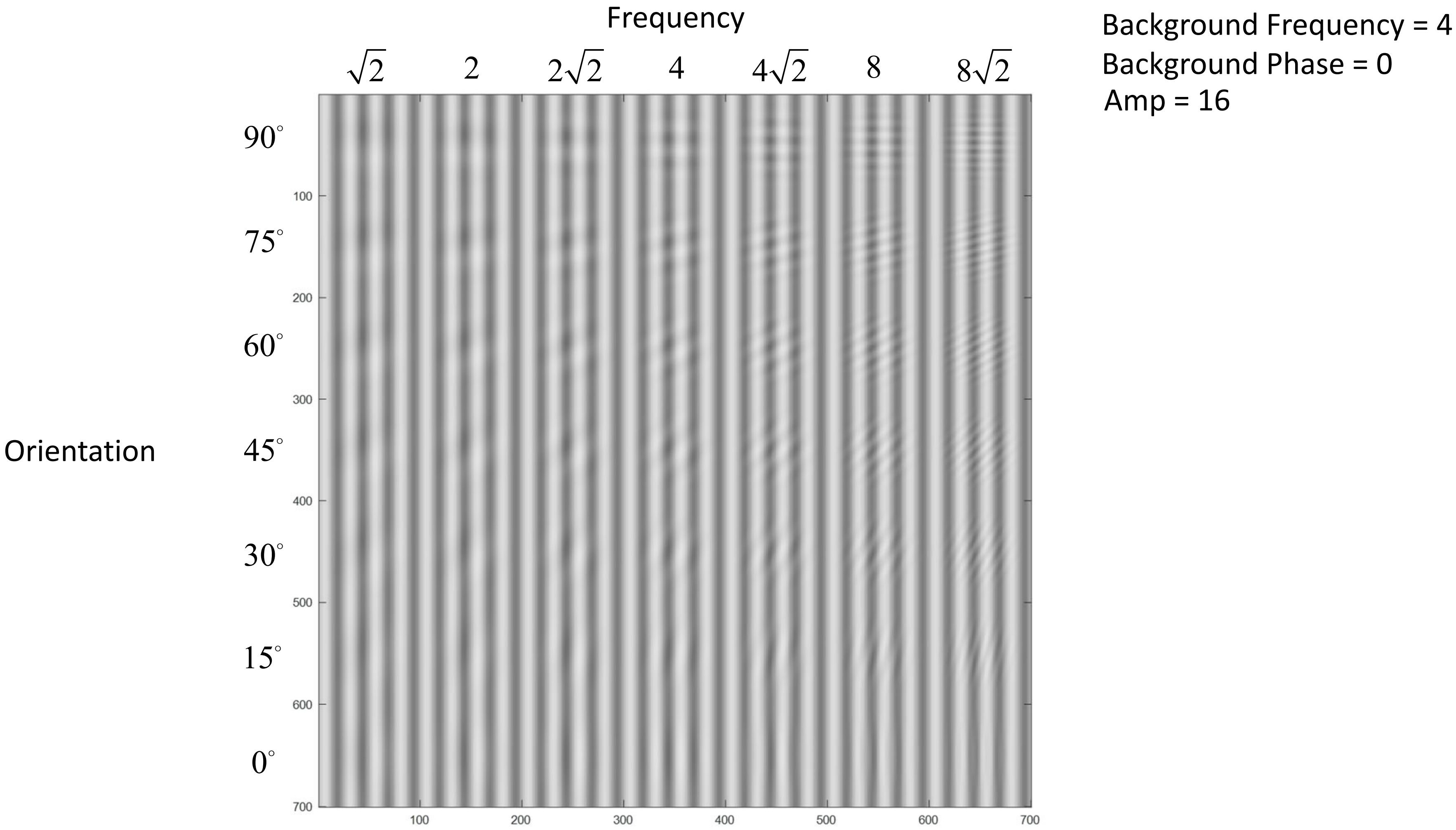
Frequency
$\sqrt{2}$
2
$2\sqrt{2}$
4
$4\sqrt{2}$
8
$8\sqrt{2}$
Orientation
$90^\circ$
$75^\circ$
$60^\circ$
$45^\circ$
$30^\circ$
$15^\circ$
$0^\circ$
100
200
300
400
500
600
700
Background Frequency = 4
Background Phase = 0
Amp = 16

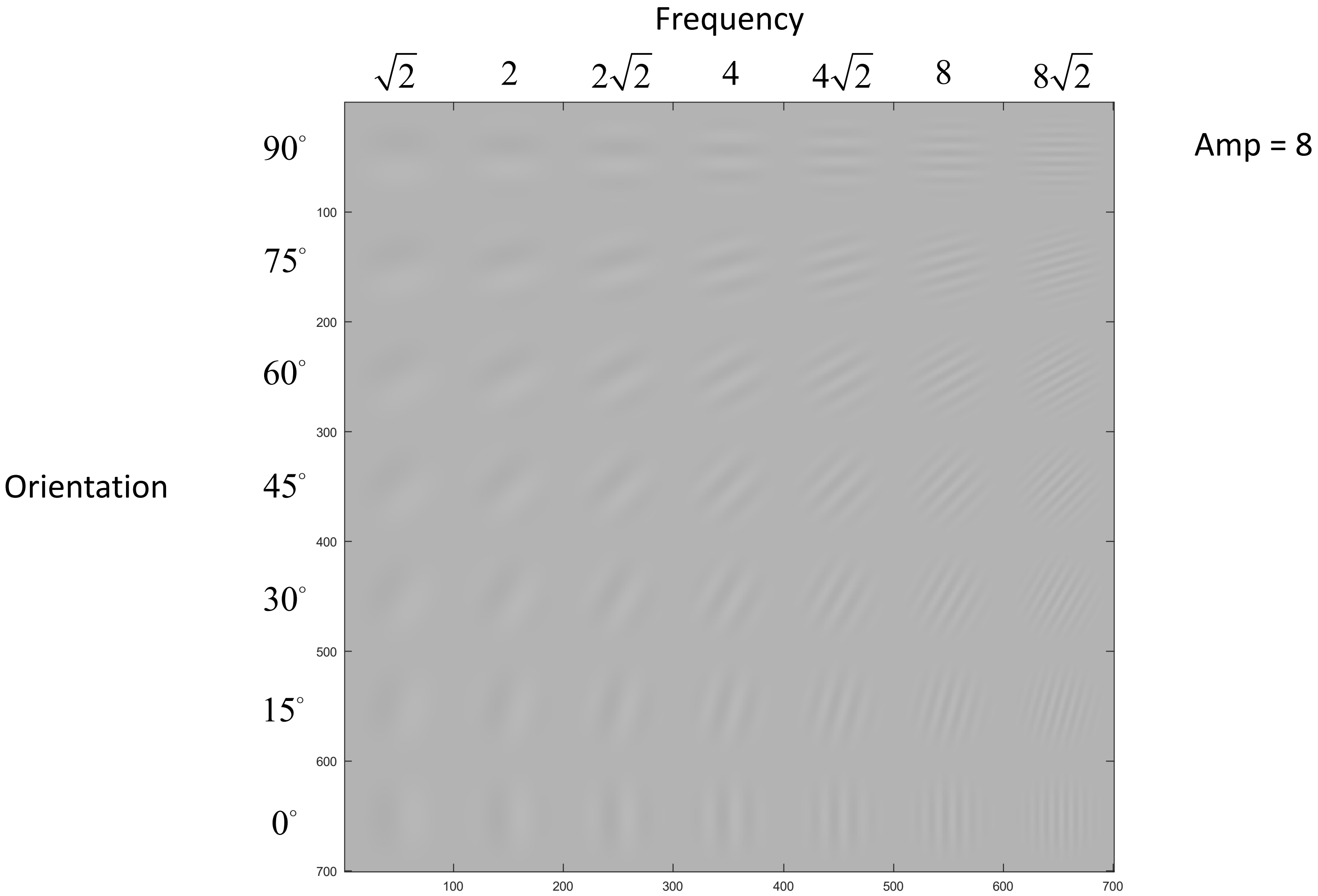
Frequency
$\sqrt{2}$
2
$2\sqrt{2}$
4
$4\sqrt{2}$
8
$8\sqrt{2}$
Amp = 8
Orientation
$90^\circ$
$75^\circ$
$60^\circ$
$45^\circ$
$30^\circ$
$15^\circ$
$0^\circ$
100
200
300
400
500
600
700

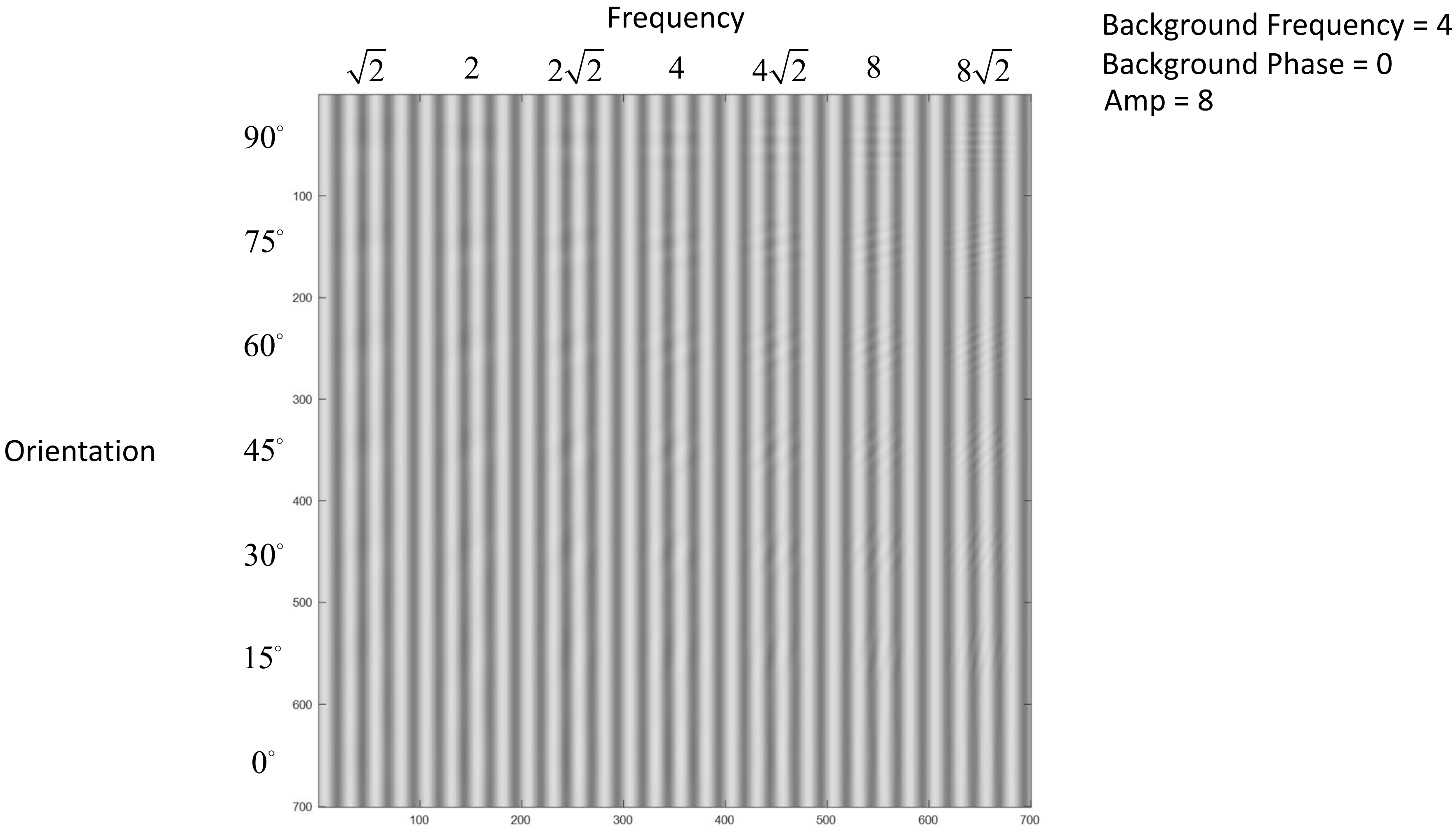

Frequency
$\sqrt{2}$
2
$2\sqrt{2}$
4
$4\sqrt{2}$
8
$8\sqrt{2}$
Orientation
$90^\circ$
$75^\circ$
$60^\circ$
$45^\circ$
$30^\circ$
$15^\circ$
$0^\circ$
100
200
300
400
500
600
700
Background Frequency = 4
Background Phase = 0
Amp = 8

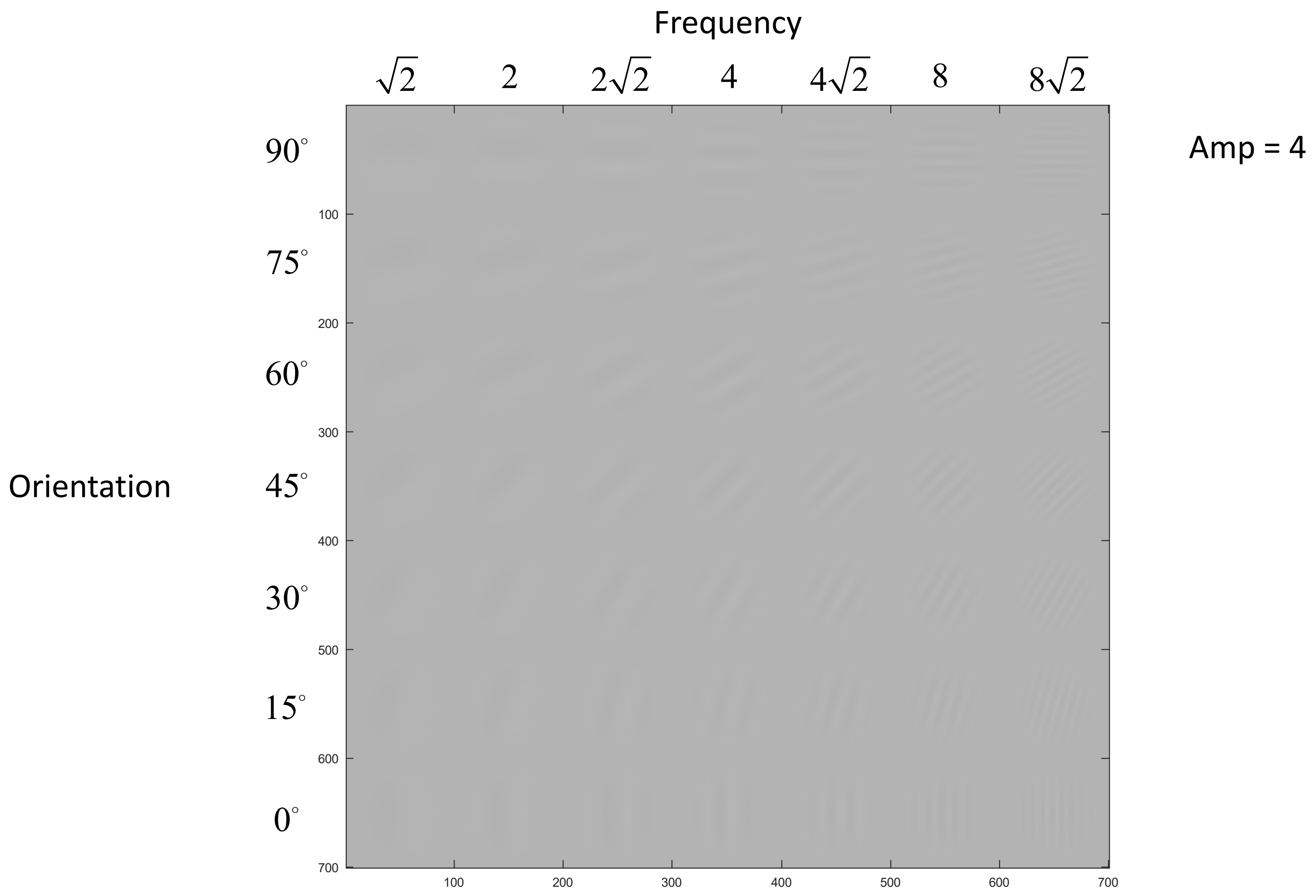

Frequency
$\sqrt{2}$
2
$2\sqrt{2}$
4
$4\sqrt{2}$
8
$8\sqrt{2}$
Amp = 4
Orientation
$90^\circ$
$75^\circ$
$60^\circ$
$45^\circ$
$30^\circ$
$15^\circ$
$0^\circ$
100
200
300
400
500
600
700

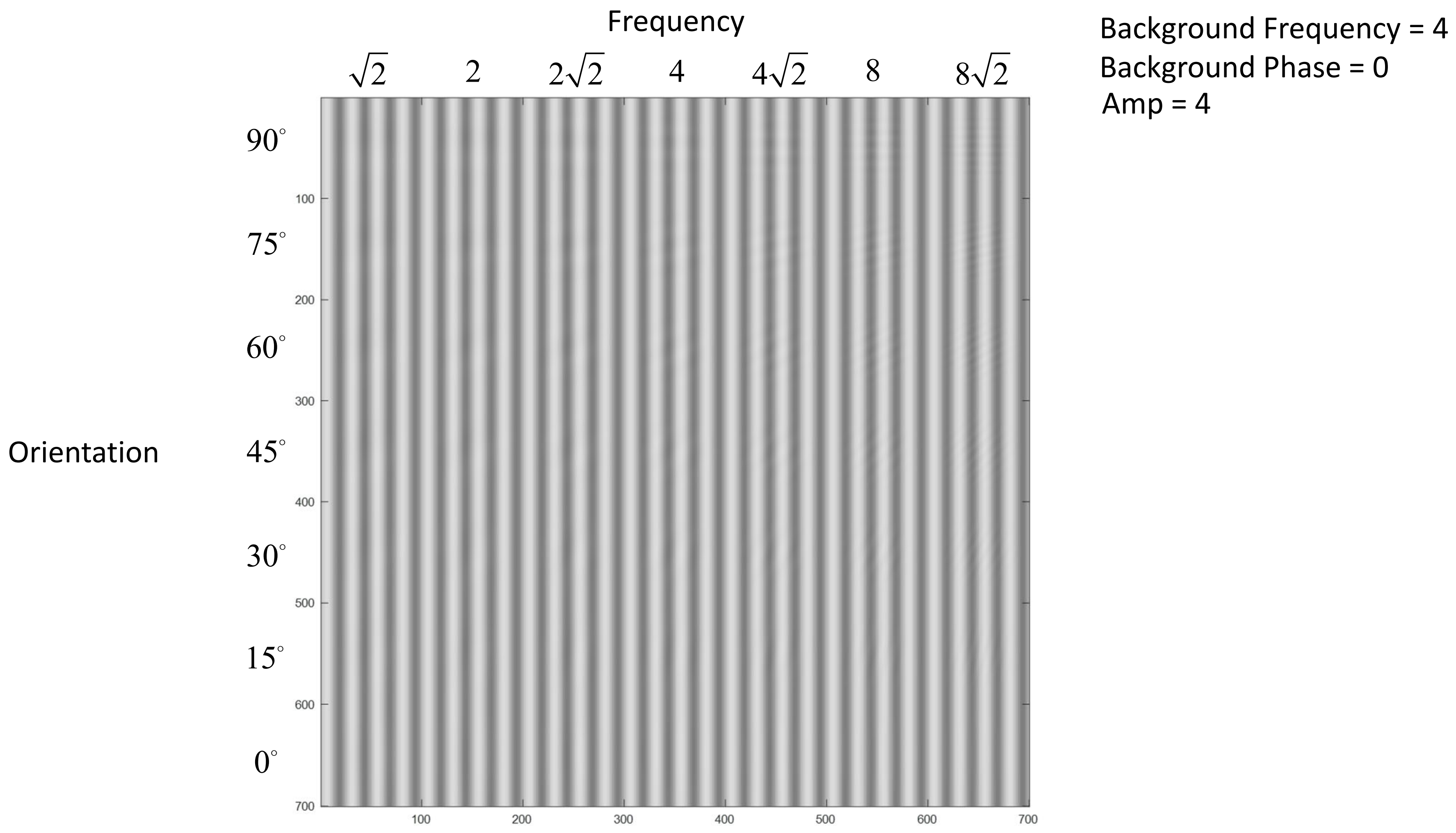
Frequency
$\sqrt{2}$
2
$2\sqrt{2}$
4
$4\sqrt{2}$
8
$8\sqrt{2}$
Orientation
$90^\circ$
$75^\circ$
$60^\circ$
$45^\circ$
$30^\circ$
$15^\circ$
$0^\circ$
100
200
300
400
500
600
700
Background Frequency = 4
Background Phase = 0
Amp = 4
